# HSV-1 single cell analysis reveals anti-viral and developmental programs activation in distinct sub-populations

**DOI:** 10.1101/566489

**Authors:** Nir Drayman, Parthiv Patel, Luke Vistain, Savaş Tay

## Abstract

Viral infection is usually studied at the population level by averaging over millions of cells. However, infection at the single-cell level is highly heterogeneous. Here, we combine live-cell imaging and single-cell RNA sequencing to characterize viral and host transcriptional heterogeneity during HSV-1 infection of primary human cells. We find extreme variability in the level of viral gene expression among individually infected cells and show that they cluster into transcriptionally distinct sub-populations. We find that anti-viral signaling is initiated in a rare group of abortively infected cells, while highly infected cells undergo cellular reprogramming to an embryonic-like transcriptional state. This reprogramming involves the recruitment of beta-catenin to the host nucleus and viral replication compartments and is required for late viral gene expression and progeny production. These findings uncover the transcriptional differences in cells with variable infection outcomes and shed new light on the manipulation of host pathways by HSV-1.

## INTRODUCTION

Viruses are obligatory intracellular parasites that rely on the biochemical functions of their hosts to carry out infection. While usually studied at the level of cell populations, viral infection is inherently a single-cell problem, where the outcome of infection can dramatically differ between genetically identical cells. For example, early studies in the 1940s investigated the burst size of individually infected bacteria and concluded that it both spans three orders of magnitude and cannot be solely attributed to differences in bacteria size (Delbrück, 1945). A later study measured the burst size from individual HeLa cells infected with Herpes Simplex virus 1 (HSV-1) and found that many of the infected cells did not release any progeny, that the variability between individual cells was high and that it did not correlate with the multiplicity of infection (MOI) used (Wildy et al., 1959). More recently, technological improvements have allowed the quantification of burst sizes and infection kinetics of different mammalian viruses, pointing to a high degree of cell-to-cell variability in infection (Zhu et al., 2009; Timm and Yin, 2012; Schulte and Andino, 2014; Combe et al., 2015; Heldt et al., 2015; Cohen and Kobiler, 2016; Guo et al., 2017; Drayman et al., 2017). One well-known source of this variability is the random distribution of the number of viruses that individual cells encounter (Parker, 1938; Smith, 1968; Cohen and Kobiler, 2016). Another source is genetic variability in the virus population, with some virus particles being unable or less fit to establish infection (Huang and Baltimore, 1970; Lauring et al., 2013; Stern et al., 2014).

It is becoming clear that a third source of this variability is the host cell state at the time of infection (Snijder et al., 2009, 2012; Drayman et al., 2017). Variability in the host cell state can arise from both deterministic processes such as the cell-cycle and stochastic processes such as mRNA transcription and protein translation (Elowitz et al., 2002; Cohen et al., 2008; Tay et al., 2010; Loewer and Lahav, 2011; Kellogg and Tay, 2015). Recently, the advent of single-cell RNA-sequencing (scRNA-seq) has allowed researchers to examine virus-host interactions in multiple systems, mainly those of RNA viruses (Steuerman et al., 2018; Russell et al., 2018; Xin et al., 2018; Zanini et al., 2018; Shnayder et al., 2018). While scRNA-seq is providing a wealth of new information on viral infection, it is currently limited to the characterization of highly abundant transcripts. Thus, it is clear that a better understanding of viral infection requires studies at the single-cell level.

HSV-1 is a common human pathogen that belongs to the *herpesviridae* family and serves as the prototypic virus for studying alpha herpesviruses infection. HSV-1 *de novo* infection has both lytic and latent phases. In the lytic phase, the virus infects epithelial cells at the site of contact, where it replicates, destroys the host cell and releases viral progeny. The latent phase is restricted to neurons, in which the virus remains silent throughout the host life with occasional reactivation. Here, we focus on the lytic part of the virus life cycle. While lytic infection is usually asymptomatic, in some cases - particularly in immune-compromised individuals and infants - it can results in life threatening conditions such as meningitis and encephalitis.

To initiate infection, HSV-1 must bind to its receptors, enter the cytoplasm, travel to the nuclear pore and inject its linear double-stranded DNA into the host nucleus (Kobiler et al., 2012). Once in the nucleus, viral gene expression proceeds in a temporal cascade of three classes of viral genes: immediate-early, early and late (Honess and Roizman, 1974, 1975; Harkness et al., 2014). DNA replication occurs in sub-nuclear structures, called replication compartments (RCs), that aggregate the seven essential replication proteins as well as other viral and host proteins (de Bruyn Kops and Knipe, 1988; Liptak et al., 1996; Weller and Coen, 2012; Dembowski and DeLuca, 2015; Dembowski et al., 2017; Reyes et al., 2017; Dembowski and DeLuca, 2018). ICP4 is the major viral trans-activator and is required for viral infection to progress beyond the point of immediate-early gene expression. Upon viral DNA replication, ICP4 is predominantly localized in the RCs, with some diffuse nuclear and cytoplasmic localization (Knipe et al., 1987; Zhu and Schaffer, 1995).

Several studies applied high-throughput technologies to analyze the cellular response to HSV-1 infection at the population level. RNA sequencing revealed a widespread deregulation of host transcription, including the disruption of transcription termination (Rutkowski et al., 2015; Hennig et al., 2018), activation of anti-sense transcription (Wyler et al., 2017), depletion of RNA-polymerase II from the majority of host genes (Abrisch et al., 2015; Birkenheuer et al., 2018) and changes in splicing and polyadenylation (Hu et al., 2016). While most cellular genes are down-regulated by infection, some genes were reported to be up-regulated, including host transcription factors and anti-viral genes (Pasieka et al., 2006; Taddeo et al., 2002; Hu et al., 2016). Proteomics studies have defined the different stages and protein complexes during HSV-1 replication (Dembowski and DeLuca, 2015; Suk and Knipe, 2015; Dembowski et al., 2017; Reyes et al., 2017; Dembowski and DeLuca, 2018) as well as the cellular protein response to infection (Kulej et al., 2017; Lum et al., 2018).

While incredibly informative, population-level analyses suffer in that they average over all the cells in the population. In the case of virus-infected cells, the population is far from homogenous and could in fact contain opposite phenotypes such as highly-infected and abortively-infected cells, leading to contradictory results. One such example is the seemingly complex relation between HSV-1 infection and type I interferon (IFN) signaling. The picture that emerges from population-level measurements is oxymoronic, with wild-type HSV-1 infection both clearly activating (Gianni et al., 2013; Hu et al., 2016; Liu et al., 2016; Reinert et al., 2016) and clearly repressing (Lin et al., 2004; Johnson et al., 2008; Kew et al., 2013; Johnson and Knipe, 2010; Su et al., 2016; Christensen et al., 2016; Manivanh et al., 2017; Yuan et al., 2018; Chiang et al., 2018) the type I IFN pathway. Such discrepancies might be resolved with the use of single-cell measurements.

Here, we apply a combination of live-cell time-lapse fluorescent imaging, scRNA-seq and sequencing of sorted cell populations to explore HSV-1 infection at the single-cell level. We find that single cells infected by the virus show variability in all aspects of infection, starting from the initial phenotype (abortive infection vs. successful initiation of viral gene expression), through the timing and rate of viral gene expression and ending with the host cellular response. This study resolves the apparent discrepancy in the literature regarding type I IFN induction and shows that it is restricted to a rare sub-population of abortively-infected cells. Surprisingly, we find that the main transcriptional response in highly-infected cells is the reprogramming of the cell to an embryonic-like state. We focus on the viral activation of the WNT/β-catenin pathway and find that β-catenin is recruited to cell nucleus and the viral RCs and is required for viral gene expression and progeny production. In addition to increasing our understanding of the basic nature of viral infection, these findings could potentially have clinical implications for the use of oncolytic HSV-1 based viruses for cancer treatment.

## RESULTS

### Viral infection dynamics varies among individual cells

We began by studying the temporal variability in viral gene expression initiation. To do so, we employed a wild-type HSV-1 (strain 17) that was genetically modified to express ICP4-YFP (Everett et al., 2003). Primary human fibroblasts (HDFn) were infected at a multiplicity of infection (MOI) of 2 and monitored by time-lapse fluorescent microscopy (Fig. 1A, supplementary Movie 1). An MOI of 2 was chosen as it resulted in ∼50% of the cells becoming ICP4-positive during primary infection. Note that we determined the genome:PFU ratio for our viral stock and found it to be 36±4, suggesting that all the cells in the culture have likely encountered numerous virus particles.

**Figure 1.**
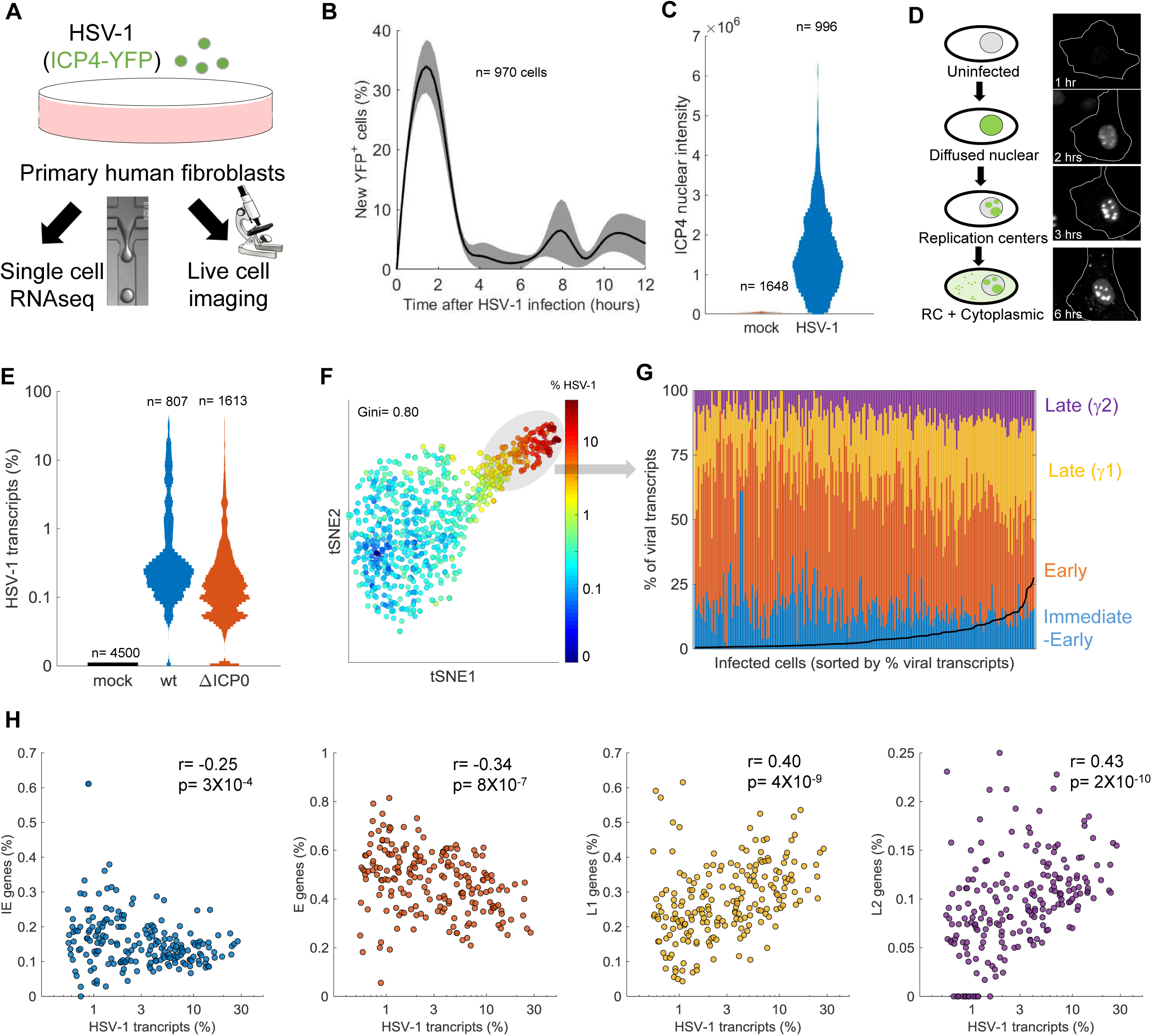
Cell-to-cell variability in infection dynamics and viral gene expression. **A.** HDFn cells were infected with HSV-1 expressing ICP4-YFP and analyzed by time-lapse fluorescent imaging and scRNA-seq. **B.** Distribution of the initial time of ICP4 expression. The black line denotes the average and the gray shadowing denotes the standard error of the time of ICP4 expression by single cells. n=970 cells from 3 fields of view. **C.** Violin plots showing the distribution of ICP4 nuclear intensity at 5 hours post infection of mock (n=1648) or HSV-1 infected (n=996) cells. **D.** Schematic diagram of the different localization phenotypes of ICP4 (left) and a representative single cell showing these phenotypes at different time points following infection. **E.** Violin plots showing the distribution of the % of viral transcripts (out of the total transcripts) for individual cells which were mock-infected (n=4500), infected with wt (n=807) or ΔICP0 (n=1613) HSV-1. Y-axis is logarithmic. **F.** tSNE plot based on viral gene expression in wt HSV-1 infection. Each dot represents a single cell and is colored according to the % of viral transcripts from blue (low) to red (high). Color bar is logarithmic. Gini is the Gini coefficient **G.** The relative abundance of the four viral gene classes: immediate-early (blue), early (orange), late γ1 (yellow) and late γ2 (purple) in highly-infected single cells (highlighted by a gray oval in panel F). Single cells are ordered by their % viral transcripts from low (left) to high (right) which is denoted by the black line. **H.** Scatter plots of single cells showing the % viral transcripts on the x-axis and the relative abundance of each viral gene class on the Y-axis. r and p are the Pearson correlation coefficients and p-values, respectively.

Initiation of ICP4 expression was observed to mostly occur between 1 and 4 hours post-infection (Fig. 1B). Almost no new infections were observed between 4-6 hours, but two infection peaks were later seen at 8 and 11 hours. These peaks are likely the result of secondary infections, since new viral progeny can be detected in infected cells starting at 6 hours post-infection (Pomeranz and Blaho, 2000; Ikeda et al., 2011; Drayman et al., 2017). Given that the majority of infected cells have initiated viral gene expression by 5 hours, we chose this time point for further analyses. At 5 hours, two cellular populations can be clearly distinguished: cells which successfully initiated viral gene expression (ICP4^+^) and cells in which infection was aborted (ICP4^-^). Of 1,814 cells infected with HSV-1, 996 cells (55%) were ICP4^+^ and 818 (45%) were ICP4^-^.

Among the ICP4^+^ cells, nuclear levels of ICP4 varied by ∼100-fold, ranging from 7×10^4^ to 9×10^6^ AU (Fig. 1C). Infected cells showed three distinct phenotypes in regards to ICP4 localization (Fig. 1D). Upon its expression, ICP4 is initially diffuse throughout the cell nucleus. As its level increases, ICP4 forms discrete foci in the nucleus. These are the viral RCs, where viral DNA replication takes place. Later, the levels of cytoplasmic ICP4 increases and interspersed foci can be seen in the cytoplasm. These phenotypes are temporally linked and delineate the progression through infection. As evident by time-lapse microscopy (supplementary Movie 1), individual cells show a high degree of variability not only in the timing of initial gene expression, but also in the rate of infection progression.

Taken together, we find that not all infected cells successfully initiate viral gene expression under these experimental conditions. Those that do initiate viral gene expression show variation in the timing of initial gene expression, the rate of infection progression and the level and localization of the immediate-early protein ICP4. These results prompted us to explore cellular heterogeneity on a larger scale by applying single-cell RNA-sequencing (scRNA-seq) to infected cells.

### Viral gene expression is extremely variable among individual cells

HDFn were mock-infected or infected with wild-type or a ΔICP0 HSV-1 mutant and harvested for scRNA-seq at 5 hours post-infection. We chose to include the ΔICP0 mutant as it results in a relatively high number of abortive infections and a robust activation of anti-viral responses. For scRNA-seq we applied the Drop-seq protocol (see Methods and (Macosko et al., 2015). Briefly, a microfluidic device was used to encapsulate individual cells in a water-in-oil droplet in which cell lysis, mRNA-capture and barcoding took place. The barcoded mRNA was then recovered from the droplets, reverse-transcribed, amplified and sequenced. Since each cDNA was barcoded with a cell and transcript ID, the sequencing data allows reliable quantification of the number of transcripts in individual cells.

Only 0.4% of mock-infected cells had any reads aligned to the HSV-1 genome, with a maximal expression of 2 viral gene counts (0.05% of transcripts). Cells infected with either wt or ΔICP0 HSV-1 showed extreme cell-to-cell variability in the amount of viral transcripts they express, ranging from 0-36% (Fig. 1E). The viral gene expression distribution was highly skewed, with most cells expressing low levels of viral transcripts and some cells expressing much higher levels (Fig. 1E). The Gini coefficient, a measurement of population inequality ranging from zero (complete equality) to one (complete inequality), was used to evaluate the distribution of viral gene expression among individual cells. The Gini coefficients were 0.8 for wt infection and 0.77 for ΔICP0, higher than that reported for viral gene expression by Influenza virus (0.64, (Russell et al., 2018)). When wt viral gene expression is visualized in two-dimension (using the tSNE dimensionality reduction technique (Maaten and Hinton, 2008)), two clusters of cells can be seen, distinguished by the amount of viral gene expression (less or more than ∼1%, Fig. 1F). A similar distribution was seen for ΔICP0 infected cells, although there were significantly less cells in the “highly-infected” cluster (Supplementary Fig. 1).

To further explore cell-to-cell variability in viral gene expression, we analyzed the relative expression of the four groups of viral transcripts, corresponding to their temporal order of expression: immediate-early (IE), early (E) and late (subdivided into early-late (γ1) and true-late (γ2)). We focused on the group of highly infected cells, since the lowly infected cells had too few viral gene counts for accurate analysis. Fig. 1G shows the relative expression of the viral gene classes in single cells, ordered from low to high viral gene expression. The fraction of late genes increases as total viral gene expression increases, at the expense of IE and E genes. The correlations between viral gene expression and the four classes of viral transcripts are shown in Fig. 1H-K. Similar observations were made for ΔICP0 infected cells (Supplementary Fig. 1).

Our scRNA-seq data indicates a wide and uneven distribution of viral gene expression during HSV-1 infection, with most cells expressing none or low levels of viral gene transcripts and a smaller group expressing much higher levels, in agreement with the ICP4 expression levels presented above. We note that significant cell-to-cell differences are seen even within the group of highly infected cells, with viral gene expression ranging from 1% to >30%, and that this “viral expression load” is correlated with late gene expression.

### The cell-cycle affects HSV-1 gene expression

The effect of the cell cycle on HSV-1 infection was evaluated by calculating a cell-cycle score for each cell in our dataset (Tirosh et al., 2016) and measuring the correlation between HSV-1 gene expression and the cell-cycle score (Supplementary Fig. 2). We found that viral gene expression is negatively correlated with the cell-cycle score, with cells in the later parts of the cell cycle expressing ∼10-fold less viral genes than those in the early part of the cycle. This finding is in agreement with previous results, showing that cells in the G2 phase of the cell-cycle are less likely to initiate viral gene expression (Drayman et al., 2017).

**Figure 2.**
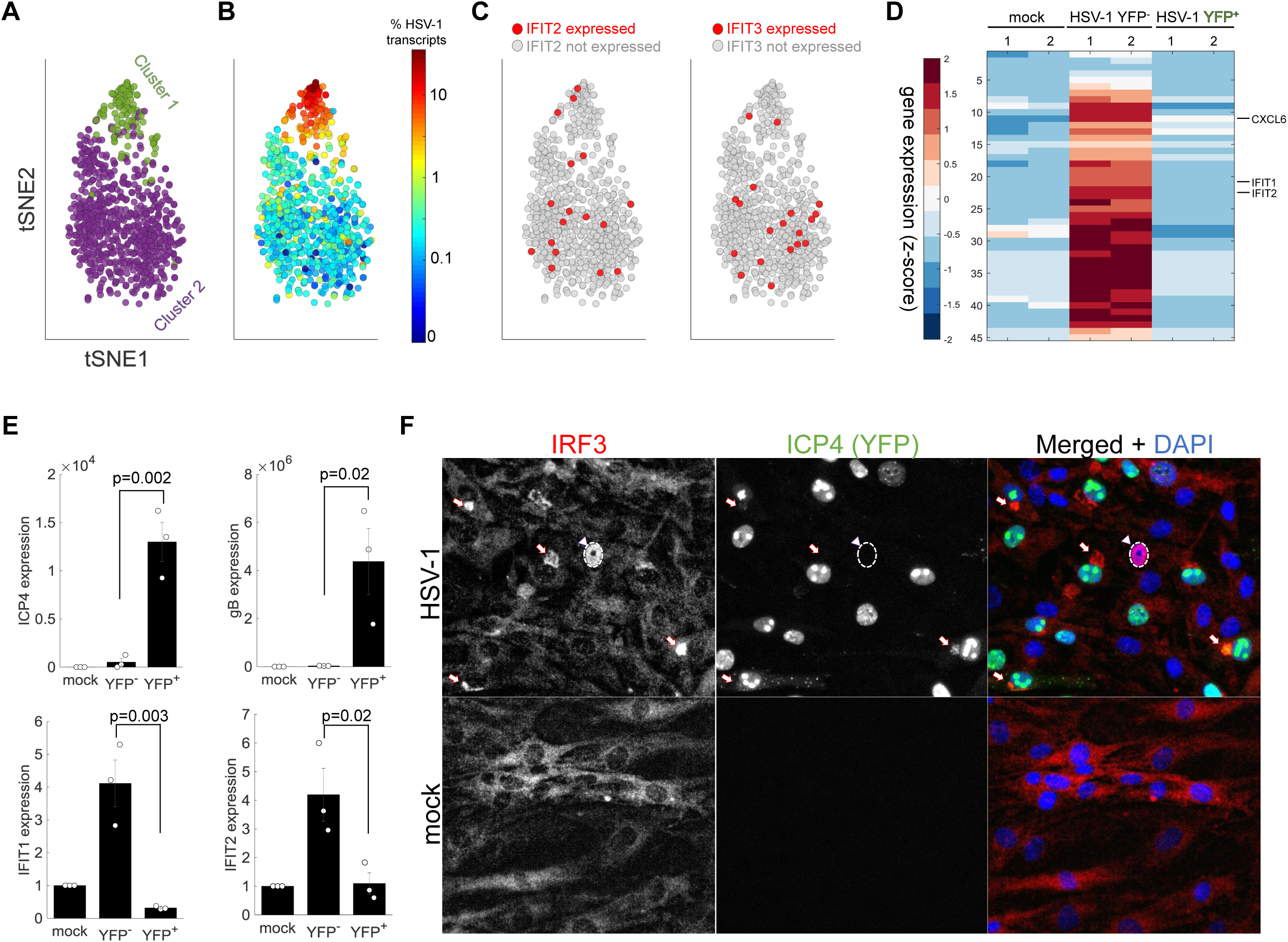
Anti-viral program is initiated in abortively infected cells. **A.** tSNE plot based on viral and host gene expression. Cells are colored according to their clustering. Cluster 1 is colored green and cluster 2 is colored purple. **B.** tSNE plot as in A, cells are colored according to their % of viral transcripts expression. **C.** tSNE plot as in A, cells are colored according to expression (red) or no expression (gray) of *IFIT2* (left) or *IFIT3* (right). **D.** Heat-map of genes which are significantly up-regulated in ICP4^-^ cells, as compared to both mock and ICP4^+^ cells. RNA-sequencing was performed in duplicates denoted by the numbers 1 and 2 on the top row. Each row shows the normalized (z-score) expression of a single gene, colored from low (blue) to high (red). **E.** QPCR validation of selected genes. Bar plots showing the expression level of the viral genes *ICP4* and *gB* (top) and the anti-viral genes *FIT1* and *IFIT2* (bottom). Value are mean±s.e of three independent biological repeats. Individual measurements are shown as circles. p-values were calculated using a one-tailed t-test **F.** Immunoflorescent staining of IRF3 in mock-infected (bottom) or HSV-1 infected (top) cells at 5 hours post infection. The arrowhead points to an ICP4 negative cell that shown nuclear IRF3 staining (nucleus border denoted by a dashed white line). The arrows point to aggregation of IRF3 outside the nucleus in ICP4 positive cells.

As the cell-cycle is both a major source of cell-to-cell variability and negatively correlated with viral gene expression, it was crucial to regress out the cell-cycle effect before analyzing the host response (Supplementary Fig. 2). We could now turn to analyze the host genes that are differentially expressed among HSV-1 infected cells, starting with the anti-viral response.

### The anti-viral program is only detected in a rare sub-population of abortively infected cells

As previous population-level studies reported the activation of anti-viral genes during wild-type HSV-1 infection, we hypothesized that highly infected cells (Fig 2A,B, cluster 1) should be enriched for anti-viral genes. To our surprise, differential gene expression analysis of the two clusters did not indicate up-regulation of the anti-viral response in cluster 1 (Sup. Table 1). In fact, canonical anti-viral genes such as *IFIT2* and *IFIT3* were only detected in 2-3% of the cells from both clusters 1 and 2 (Fig. 2C).

One possible explanation is that anti-viral genes are indeed expressed in highly infected cells but were not detected by scRNA-seq due to technical limitations. To investigate this, infected cells were FACS-sorted into two populations based on ICP4-YFP expression (ICP4^+^ and ICP4^-^) and each population was sequenced. In agreement with the scRNA-seq data, expression of canonical anti-viral genes was not significantly different between mock-infected and ICP4^+^ cells. Rather, our analysis indicated that a small group of genes, including the anti-viral genes *IFIT1* and *IFIT2*, were specifically up-regulated in the ICP4^-^population (Fig. 2D, Sup. Table 2). The Gene Ontology (GO) biological processes associated with these up-regulated genes included terms such as “response to type I interferon” and “immune response” (Sup. Table 3). QPCR validation of selected transcripts is shown in Fig. 2E.

To pinpoint the origin of the anti-viral response cells were stained for IRF3, as IRF3 translocation from the cytoplasm to the nucleus is one of the first steps in type I interferon response. In ICP4^+^ cells, IRF3 was blocked from entering the nucleus and concentrated in the nuclear periphery (Fig. 2F), while a rare subset of ICP4^-^ cells (<1%) showed nuclear localization of IRF3.

We next evaluated the anti-viral response in cells infected by ΔICP0 HSV-1. ICP0 is a multifunctional viral protein, which blocks IRF3 signaling (Lin et al., 2004). Cells infected with ΔICP0 clustered into four groups (Fig. 3A). Cluster 1 consists of abortively-infected cells with very few viral transcripts. Clusters 2 and 3 have slightly higher viral gene expression and the small cluster 4 consists of highly-infected cells (Fig. 3B,D). While the magnitude of the anti-viral response in ΔICP0-infected cells was much greater than that of wt-infected cells (Fig. 2), it was still only observed in a small population of cells, with ∼8% of the cells expressing *IFIT1* and *MX2* (compared to none of the mock-infected cells). These cells had low viral gene expression levels and belonged to clusters 1-3 (Fig. 3C,D). Anti-viral signaling was not seen in highly-infected cells of cluster 4 (Fig. 3C,D).

**Figure 3.**
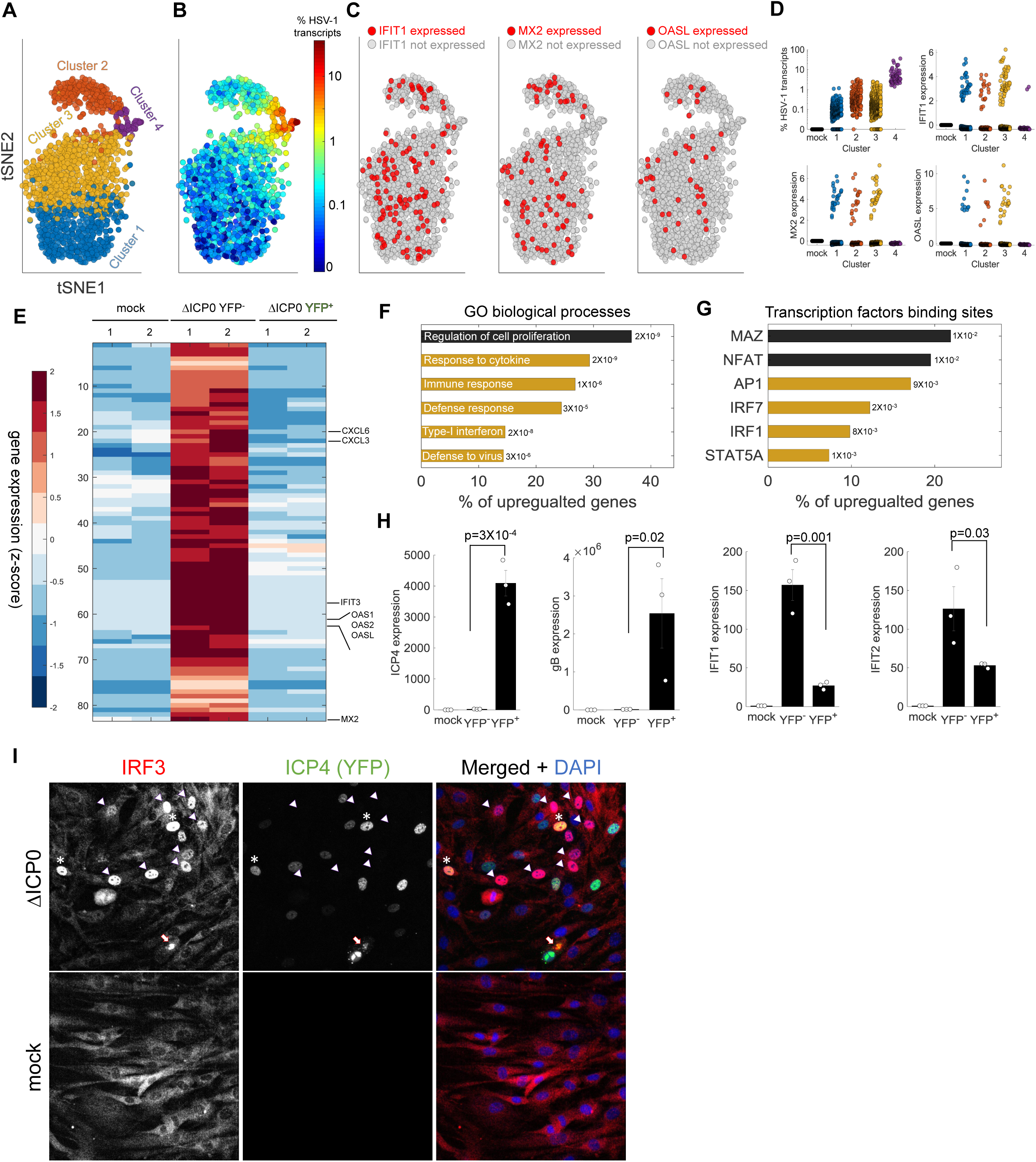
anti viral signaling in cells infected by ΔICP0 mutant. **A.** tSNE plot based on viral and host gene expression. Cells are colored according to their clustering. **B.** tSNE plot as in A, with cells colored according to the % of viral transcripts expression. **C.** tSNE plots as in A, cells are colored according to expression (red) or no expression (gray) of *IFIT1* (left), *MX2* (middle) or *OASL* (right). **D.** Scatter plots showing the expression of HSV-1 transcripts (top left) or the expression of anti-viral genes in mock-infected cells (first column) or each cluster of cells infected by ΔICP0. **E.** Heat-map of genes which are significantly up-regulated in ΔICP0-infected ICP4^-^ cells, as compared to both mock and ICP4^+^ cells. RNA-sequencing was performed in duplicates denoted by the numbers 1 and 2 on the top row. Each row shows the normalized (z-score) expression of a single gene, colored from low (blue) to high (red). **F.** GO terms enriched for genes identified in panel E. The term is written on the bar and the value next to each bar is the FDR-corrected p-value for its enrichment. Bar length shows the % of up-regulated genes annotated to the GO term. Highlighted in yellow are GO terms associated with anti-viral signaling. **G.** Transcription factor binding sites enriched in the promoters of genes identified in panel E. **H.** QPCR validation of selected genes. Bar plots showing the expression level of the viral genes *ICP4* and *gB* and the anti-viral genes *FIT1* and *IFIT2.* Value are mean±s.e of three independent biological repeats. Individual measurements are shown as circles. p-values were calculated using a one-tailed t-test **I.** Immunoflorescent staining of IRF3 in mock-infected (bottom) or DICP0 infected (top) cells at 5 hours post infection. The arrowheads point to ICP4 negative cells with nuclear IRF3. Asterisks denote ICP4 positive cells with nuclear IRF3. The arrow points to a cell in the late stages of infection, in which IRF3 is aggregated in the nuclear periphery.

RNA-sequencing of sorted cells that were infected with ΔICP0 identified ∼80 genes as significantly up-regulated in ICP4^-^ cells compared to mock and ICP4^+^ cells (Fig. 3E, Supplementary Table 4). These genes were enriched for functional annotations of anti-viral signaling (Fig. 3F, Supplementary Table 5) and binding sites of the transcription factors IRF1, IRF7 and STAT5 (Fig. 3G, Supplementary Table 6). An important difference from wt infection is that, while enriched in the ICP4^-^ population, these anti-viral genes are also activated in the ICP4^+^ population, albeit to a lesser extent (Fig. 3H). We confirmed this observation through immunofluorescent staining of IRF3 (Fig. 3I). The staining showed a higher proportion of cells with nuclear IRF3 localization compared to wt-infected cells. The majority of cells with nuclear IRF3 were ICP4^-^ but some ICP4^+^ cells also showed nuclear IRF3 staining. These ICP4^+^ cells showed diffuse nuclear localization of ICP4, which indicates that infection was aborted prior to the generation of replication compartments (Fig. 1D). The few cells that were able to proceed to the later stages of infection (as indicated by the appearance of replication compartments and cytoplasmic ICP4 foci) showed the same peri-nuclear aggregation of IRF3 as wt-infected cells (Fig. 3I and Fig. 2F).

Altogether, sequencing of both single cells and sorted cell populations, as well as immuno-fluorescence staining of IRF3, suggests that the anti-viral program is initiated in a small subset of abortively-infected cells but is blocked in highly infected cells, even in the absence of ICP0. This behavior explains the apparent discrepancy between previous population-level measurements that showed both activation and inhibition of type I interferon signaling during HSV-1 infection.

### HSV-1 infection results in transcriptional reprogramming of the host to an embryonic-like state

We next focused on genes that are up-regulated during HSV-1 infection, in either the scRNA-seq or sorted cell population experiments. 977 genes were significantly up-regulated in wt ICP4^+^ cells as compared to both ICP4^-^ and mock-infected cells (Fig. 4A, Sup. Table 7). 87 genes were significantly up-regulated in highly infected single cells (Fig. 4B, Sup. Table 1). Remarkably, we found that a major portion of these up-regulated genes are associated with GO terms that concern regulation of RNA transcription and developmental processes (Fig. 4C,D and Sup. Tables 8-9). Similar results were observed in cells infected with ΔICP0 (Supplementary Fig. 3 and Supplementary Tables 10-15).

**Figure 4.**
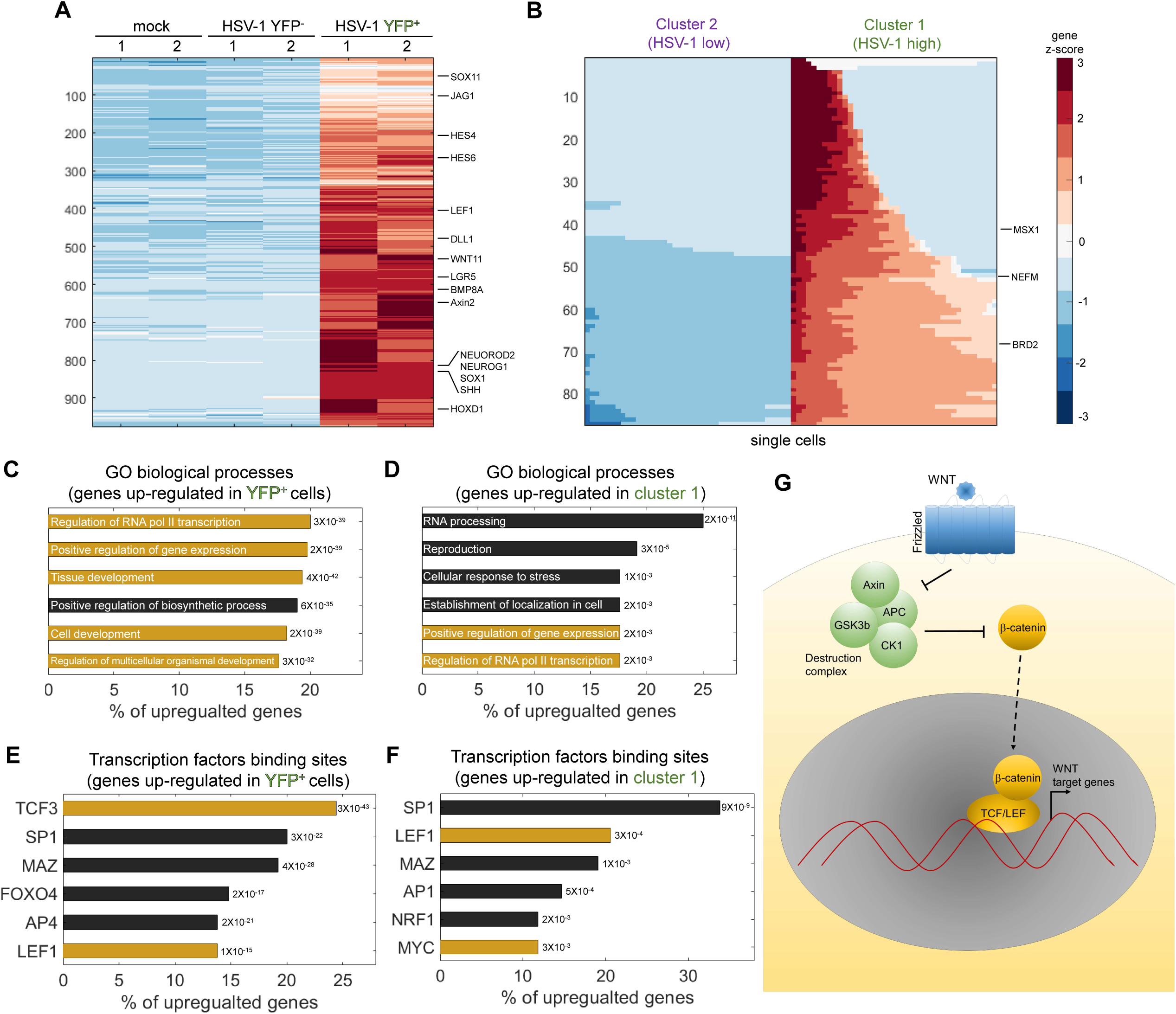
HSV-1 infection up-regulates developmental pathways. **A.** Heat-map of genes which are significantly up-regulated in ICP4^+^ cells, as compared to both mock and ICP4^-^ cells. RNA-sequencing was performed in duplicates denoted by the numbers 1 and 2 on the top row. Each row shows the normalized (z-score) expression of a single gene, colored from low (blue) to high (red). **B.** Heat-map of genes which are significantly up-regulated in cluster 1 (highly-infected) compared to cluster 2 (lowly-infected) single cell. Each row shows the normalized (z-score) expression of a single gene, colored from low (blue) to high (red) in 80 single cells (40 from cluster 1 and 40 from cluster 2). Color bar is shared for both panels A and B. **C,D.** Go terms enriched for genes identified in panels A and B, respectively. The GO term is written on the bar and the value next to each bar is the FDR-corrected p-value for its enrichment. Bar length shows the % of up-regulated genes annotated to the GO term. Highlighted in yellow are GO terms associated with development and gene regulation. **E,F.** Transcription factors binding sites enriched in the promoters of genes identified in panels A and B, respectively. The value next to each bar is the FDR-corrected p-value for its enrichment. Bar length shows the % of up-regulated genes containing binding sites for each transcription factor. Highlighted in yellow are the LEF1, TCF3 and MYC transcription factors, which are part of the WNT/β-catenin pathway. **G.** A simplified diagram of the WNT/β-catenin signaling pathway that controls LEF/TCF transcriptional activity.

The promoters of these genes are enriched for binding sites of several transcription factors, including Sp1, MAZ, LEF1 and TCF3 (Fig. 4E,F and Sup. Tables 16-17). 23% of the promoters of the up-regulated genes in ICP4^+^ cells contained a binding site for the TCF/LEF transcription factors, which are activated by the WNT/β-catenin pathway (Sokol, 2011) (Fig. 4G). Note that *LEF1* itself is a WNT target gene and is up-regulated during HSV-1 infection (Fig. 5A).

**Figure 5.**
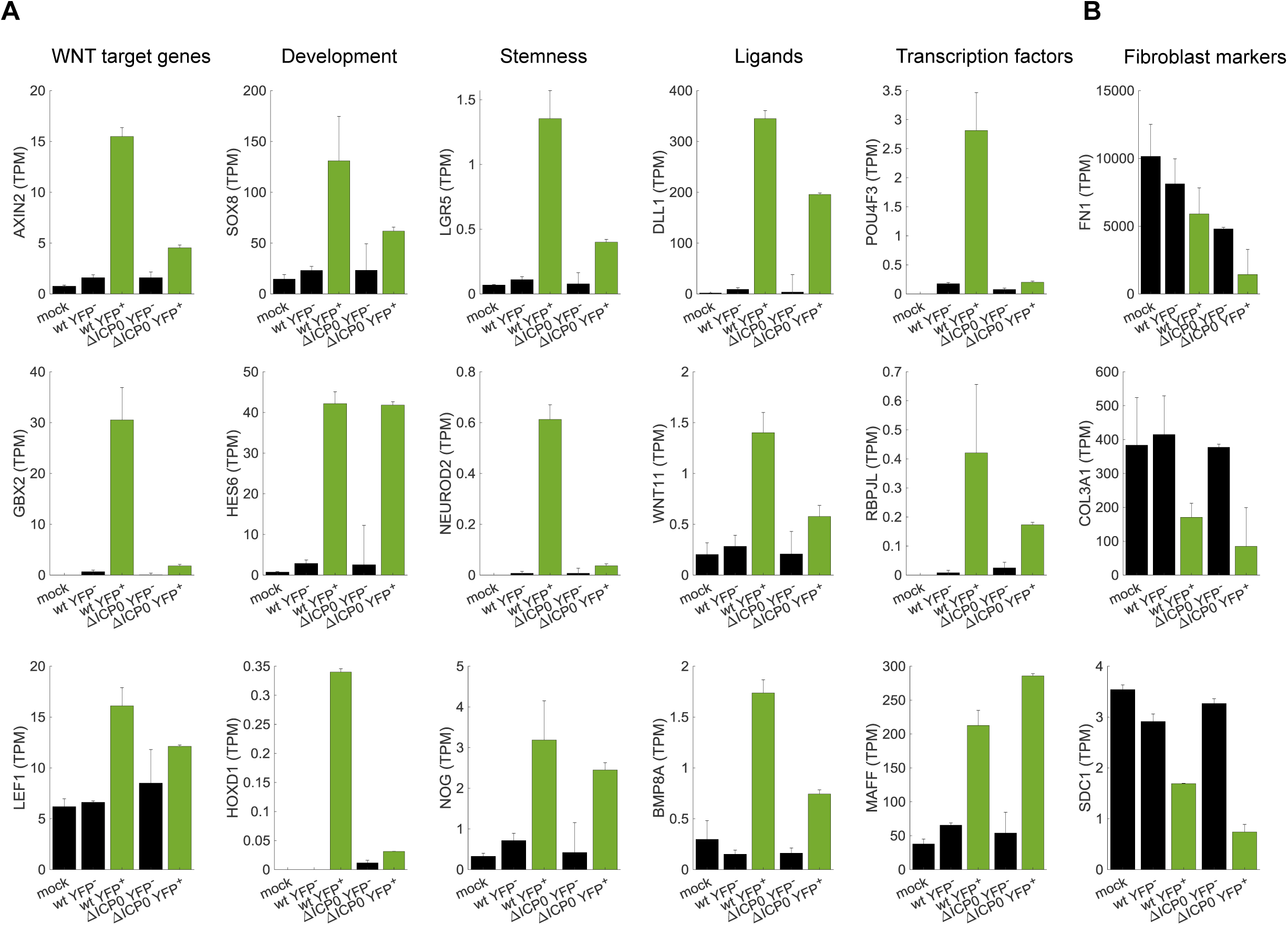
Cellular reprogramming during HSV-1 infection. **A.** Bar plots showing the expression level (TPM - transcripts per million) of selected examples of genes that participate in developmental pathways and are up-regulated in HSV-1 infected cells. Black bars denote mock-infected and ICP4^-^ cells. Green bars denote ICP4^+^ cells. Value are mean±s.e of the two sequenced biological replicates (Fig. 4A and Supplementary Fig. 3A). **B.** Bar plots showing the expression level (TPM - transcripts per million) of selected examples of fibroblast marker genes.

Examples of up-regulated genes are shown in Fig. 5A and include canonical WNT target genes such as *AXIN2*, key developmental genes belonging to the *SOX, HOX* and *HES* families, stem-cell associated transcripts such as *LGR5* and a multitude of extra-cellular ligands of various developmental pathways, including the WNT, Notch, Hedgehog and TGFβ signaling pathways. In agreement with the less efficient infection by ΔICP0, most of these transcripts are also up-regulated in ΔICP0-infected ICP4^+^ cells, but to a lesser degree than in wt-infected cells. Concomitant with the establishment of this embryonic-like transcriptional program, we observed a reduction in the levels of key fibroblast marker-genes, such as α1(III) collagen and fibronectin (Fig. 5B).

We conclude that cells highly-infected by HSV-1 undergo de-repression of embryonic and developmental genetic programs, including the WNT/β-catenin pathway.

### β-catenin translocates to the nucleus and concentrates in the viral replication compartments

Since many of the up-regulated genes are known WNT target genes and/or contain LEF/TCF binding sites in their promoters, we investigated the state of β-catenin in infected cells. Infected cells were fixed and stained for β-catenin at 5 hours post-infection (Fig. 6A). As expected, β-catenin was mainly cytoplasmic/membrane-bound in mock-infected cells. In HSV-1 infected cells, β-catenin showed three distinct localization patterns: un-perturbed (cytoplasmic), diffuse nuclear or aggregated in nuclear foci (Fig. 6A,B). Similar results were obtained for cells infected by ΔICP0 (Supplementary Fig. 4).

**Figure 6.**
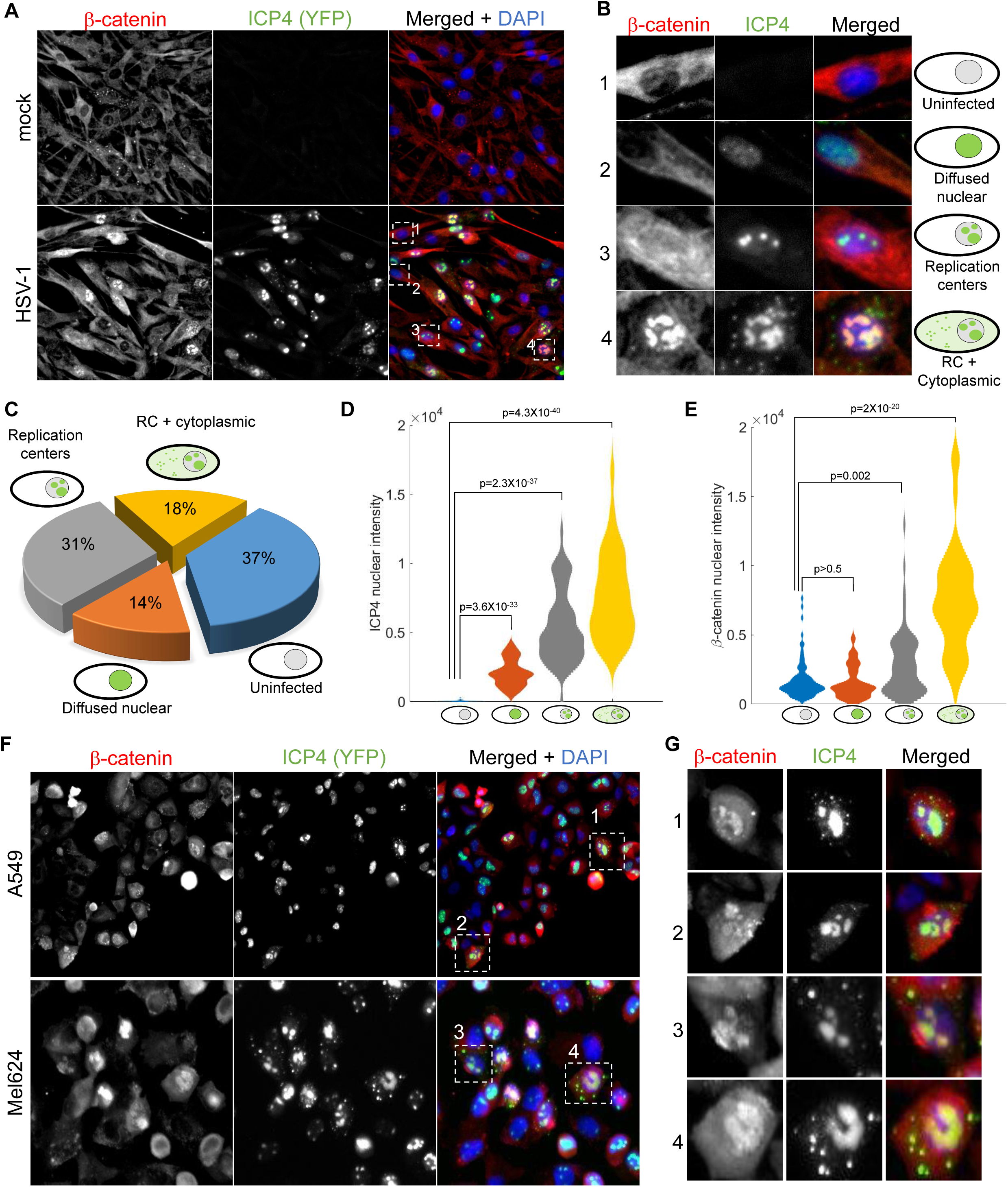
b-catenin translocates to the nucleus and concentrates in the viral replication compartments. **A.** Immunoflorescent staining of β-catenin in mock-infected (top) or HSV-1 infected (bottom) HDFn cells at 5 hours post infection. **B.** magnified imaged of the four cells denoted by dashed white boxes in panel A, showing representative images of cells with different ICP4 and β-catenin localizations. **C.** quantification of the relative abundances of the different ICP4 localizations (n=204 cells). **D,E.** Violin plots showing the distributions of ICP4 (D) and β-catenin (E) nuclear levels in cells showing the four ICP4 localization phenotypes. p-values were calculated by a two-tailed two-sample t-test and corrected for multiple comparisons by the Bonferroni correction. **F,G.** Immunofluorsence as in panels A and B for A549 and Mel624 cells.

At 5 hours post-infection, 37% of the cells were ICP4 negative, 14% were at the earliest stage of infection (diffuse nuclear ICP4), 31% have assembled viral replication compartments and 18% progressed to show cytoplasmic foci of ICP4 (Fig. 5C). ICP4 levels increase from one group to the next, in accordance with the temporal progression of infection (Fig. 6D).

β-catenin localization was linked to the temporal progression of infection. β-catenin remained cytoplasmic in both ICP4-negative cells and cells with diffused nuclear ICP4, translocated to the nucleus upon the generation of the replication compartments and subsequently co-localized with the RCs, but only in cells showing cytoplasmic foci of ICP4 (Fig. 6B,E). A similar phenotype was seen in two epithelial cell-lines: A549, a lung cancer cell-line, and Mel624, a patient-derived melanoma cell-line (Fig. 6F).

This analysis indicates that β-catenin is indeed co-opted by HSV-1. Cell-to-cell variability in the progression of infection results in heterogeneity of β-catenin localization, with recruitment of β-catenin to the viral replication compartments occurring at the later stages of infection.

### β-catenin activation is necessary for late viral gene expression and progeny production

Since β-catenin target genes are activated by the virus, and since β-catenin is recruited to the viral replication centers, we hypothesized that β-catenin activity is required for the completion of the viral life cycle. We infected cells treated with an inhibitor of β-catenin activity, iCRT14 (Gonsalves et al., 2011), and measured viral gene expression at 5 hours post-infection (Fig. 7A-D). Our results indicate that β-catenin inhibition had no or minimal impact on immediate-early gene expression (Fig. 7A) but significantly inhibited early and late gene expression (Fig. 7B-D). These observations are in agreement with the late recruitment of β-catenin to the viral RCs described above.

**Figure 7.**
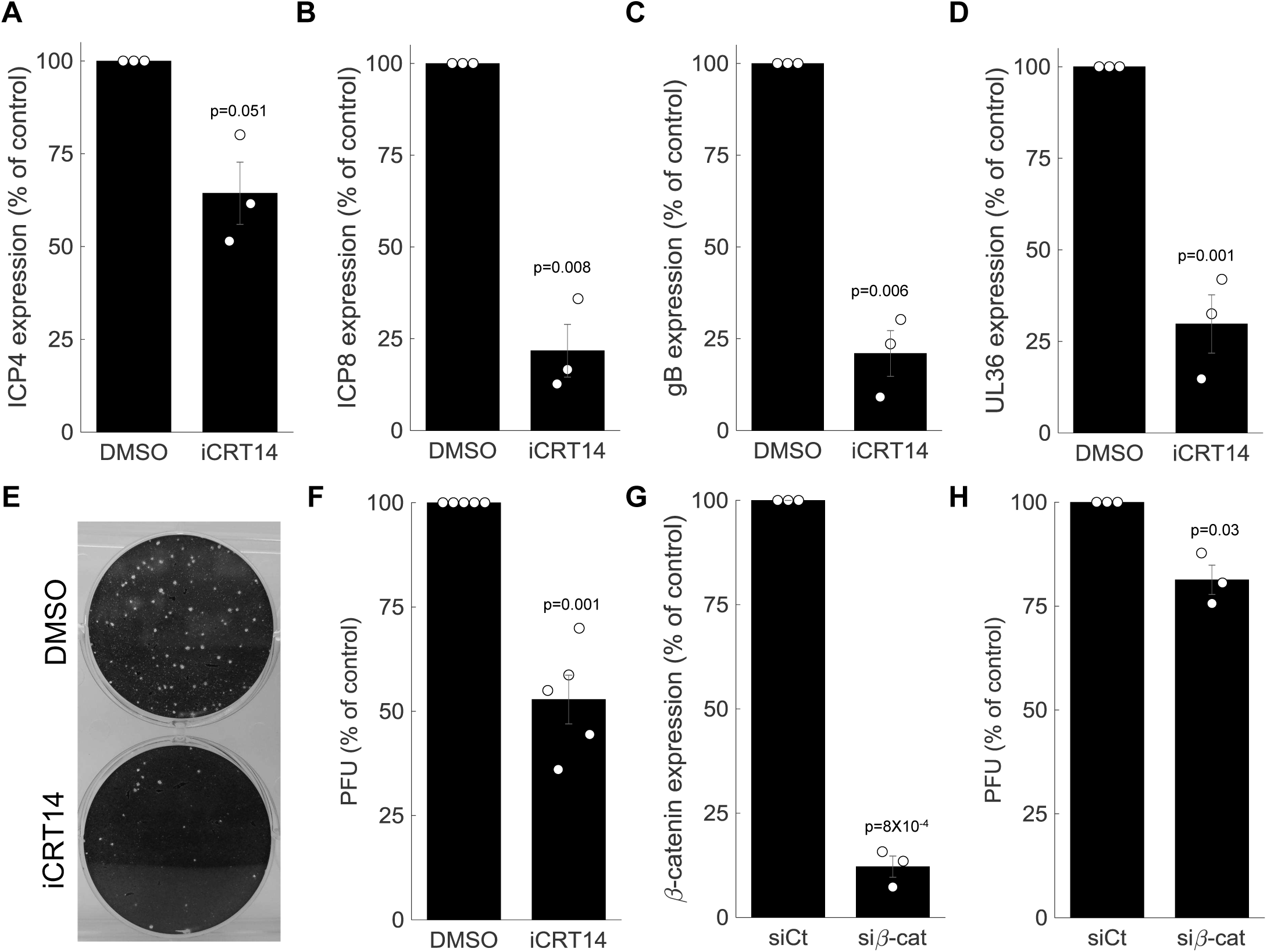
β-catenin is required for late viral gene expression and progeny production. **A-D.** HDFn treated with the β-catenin inhibitor iCRT14 or with vehicle alone (DMSO) were infected with HSV-1 and analyzed for viral gene expression by RT-PCR at 5 hours post infection. Results show the mean±standard error from three biological repeats, as compared to the DMSO treatment. The genes analyzed were ICP4 (A), ICP8 (B), gB (C) and UL36 (D). Circles are the results of individual experiments. p-values were calculated using a two-tailed one-sample t-test. **E.** Representative image of a plaque assay, titrating virus generated from a 24 hour infection of HDFn treated with iCRT14 or DMSO. **F.** Titers were calculated for viral progeny produced from iCRT14 or DMSO treated as in E. bars show the mean±standard error from five biological repeats, as compared to the DMSO treatment. Circles are the results of individual experiments. p-value was calculated using a two-tailed one-sample t-test. **G-H.** HDFn were treated with control siRNA (siCt) or siRNA targeting β-catenin (siβ-cat) for 72 hrs. Cells were assessed for β-catenin expression (G) or infected for 24 hours and viruses harvested and titrated as in panel E (H). bars show the mean±standard error from three biological repeats. p-value was calculated using a two-tailed one-sample t-test.

To measure the impact of β-catenin inhibition on viral progeny formation, we treated the cells with iCRT14, infected the cells for 24 hours and harvested and titrated the resulting viral progeny by plaque assay (Fig. 7E). In accordance with its impact on late viral gene expression, β-catenin inhibition significantly reduced viral progeny formation (Fig. 7F). Similar results were obtained when β-catenin was silenced using siRNA (Fig. 7G,H).

Taken together, our data show that HSV-1 reprograms the cell to an embryonic-like state, in part through the co-option of β-catenin, which is needed for late viral gene expression and progeny production.

## DISCUSSION

In this study we applied a combination of time-lapse fluorescent imaging, scRNA-seq and sequencing of sorted cell populations to understand HSV-1 infection at the single cell level. We find that single cells infected by the virus show variability in all aspects of infection, starting from the initial phenotype (abortive infection vs. successful initiation of viral gene expression), through the timing and rate of viral gene expression and ending with the host cellular response. Such heterogeneity in the population of infected cells is detrimental when performing population-averaged measurements but can be untangled through single-cell analyses to gain new insights into biological processes.

With regard to IFN response, we find that two opposite phenotype exist in the population of infected cells, explaining the discrepancy in the literature. Surprisingly, we find the IFN indication is limited to a small group of abortively infected cells, even in cells infected by the ΔICP0 mutant (Fig. 2,3). Why only a subset of cell activate the anti-viral program is an intriguing question and several hypotheses come to mind. For example, these cells could be poised for IFN induction due to stochastic variability in expression of the signaling pathway components (Zhao et al., 2012; Patil et al., 2015) or it might be linked to the number of viral particles a cell encounters. We plan to pursue and further characterize these rare cells in future studies.

We further found that highly-infected cells undergo transcriptional reprogramming and activate multiple developmental pathways. Focusing on β-catenin, we found it localization correspond to distinct stages in HSV-1 infection (Fig. 6): it is first recruited to the cell nucleus and then later to the viral replication compartments. These findings augment a growing body of literature that shows a link between viral infection and the β-catenin pathway (reviewed in (van Zuylen et al., 2016)), such as during infections with MCMV (Juranic Lisnic et al., 2013), Influenza (More et al., 2018), HBV (Daud et al., 2017) and Rift Valley Fever virus (Harmon et al., 2016).

How HSV-1 infection causes this massive reprogramming of the host cell state is currently unknown, although we can rule out a direct involvement of ICP0, since this reprogramming is also occurring during infection with ΔICP0, albeit to a lesser extent. While β-catenin activation is certainly one part of this, it is likely that the expression of epigenetic regulators is also important. Indeed, a proteomics study of the host chromatin during HSV-1 infection has identified widespread changes in the epigenome landscape of infected cells (Kulej et al., 2017). While we describe here a positive role for β-catenin activation during HSV-1 infection, a recent report described an inhibitory role for the germline transcription factor DUX4 during HSV-1 infection (Full et al., 2018). Thus, future studies will need to tease apart the differential contributions and effects of different developmental pathways activated during infection.

β-catenin and other developmental pathways are often found to be mutated or dysregulated during tumorigenesis (Reya and Clevers, 2005; Krausova and Korinek, 2014; Duchartre et al., 2016) and are considered as promising targets for cancer treatment (Takebe et al., 2015). In this context, it is interesting to speculate as to the possible impact of β-catenin activation by HSV-1 on its use as an oncolytic agent (Sanchala et al., 2017; Watanabe and Goshima, 2018). Currently, the first-line treatment for late-stage melanoma is immune checkpoint inhibitors (Tracey and Vij, 2019). While these inhibitors revolutionized melanoma treatment, not all patients respond to them. This was shown to be associated with β-catenin activity in the tumor, where high β-catenin levels negatively correlate with treatment success (Spranger and Gajewski, 2015; Spranger et al., 2015). Given that an HSV-1 based oncolytic therapy has been FDA approved for late-stage melanoma (Pol et al., 2015), it is tempting to speculate that the high level of β- catenin in melanomas that are resistant to checkpoint inhibitors would serve to augment oncolytic HSV-1 replication and anti-tumor effects - although this of course would have to be carefully assessed in separate studies.

**Supplementary Figure 1.**
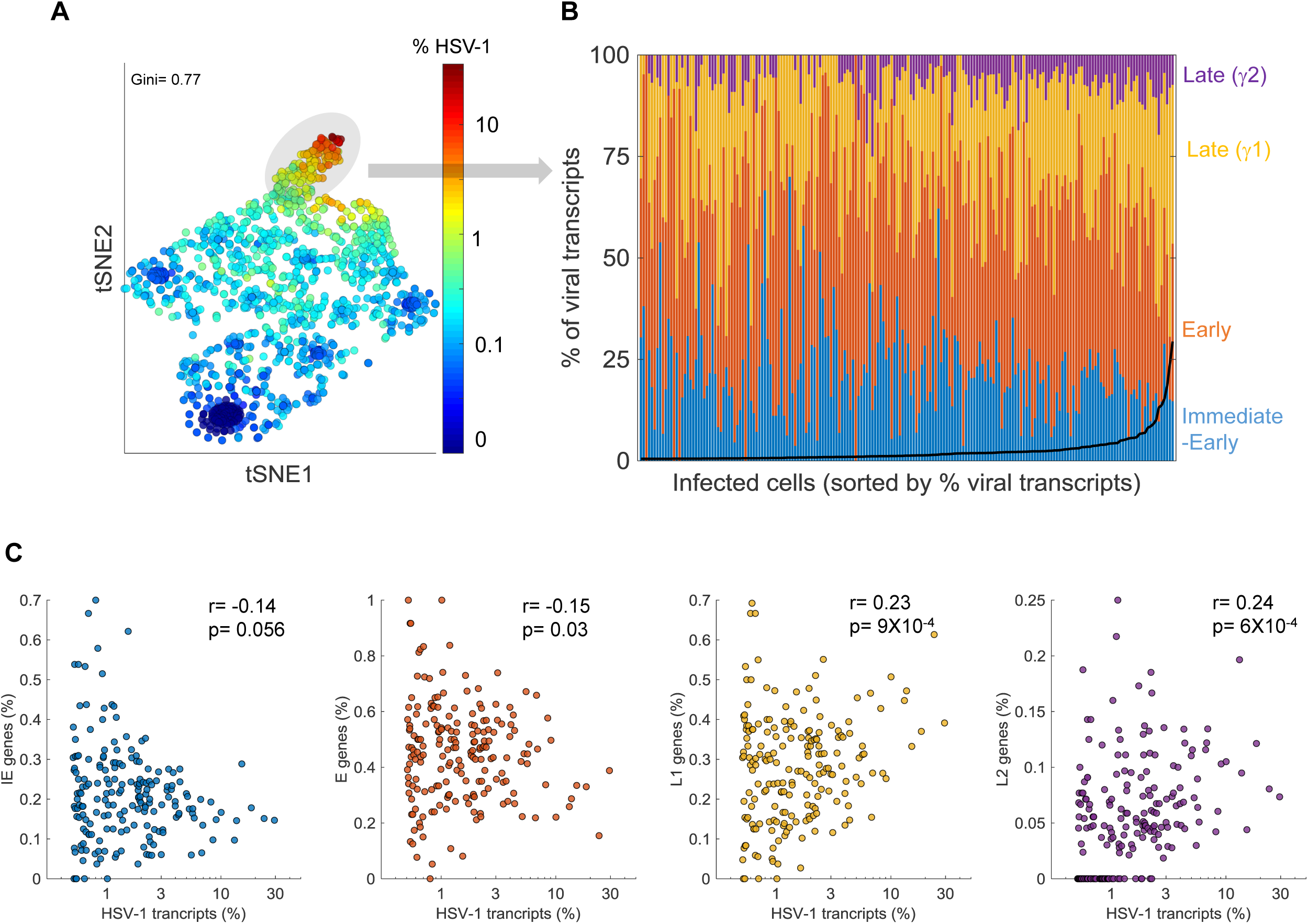
Cell-to-cell variability in viral gene expression upon ΔICP0 infection. **A** tSNE plot based on viral gene expression. Each dot represents a single cell and is colored according to the % of viral transcripts from blue (low) to red (high). Color bar is logarithmic. Gini is the Gini coefficient **B.** The relative abundance of the four viral gene classes: immediate-early (blue), early (orange), late γ1 (yellow) and late γ2 (purple) in highly-infected single cells (highlighted by a gray oval in panel A). Single cells are ordered by their % viral transcripts from low (left) to high (right) which is denoted by the black line. **C.** Scatter plots of single cells showing the % viral transcripts on the x-axis and the relative abundance of each viral gene class on the Y-axis. r and p are the Pearson correlation coefficients and p-values, respectively.

**Supplementary Figure 2.**
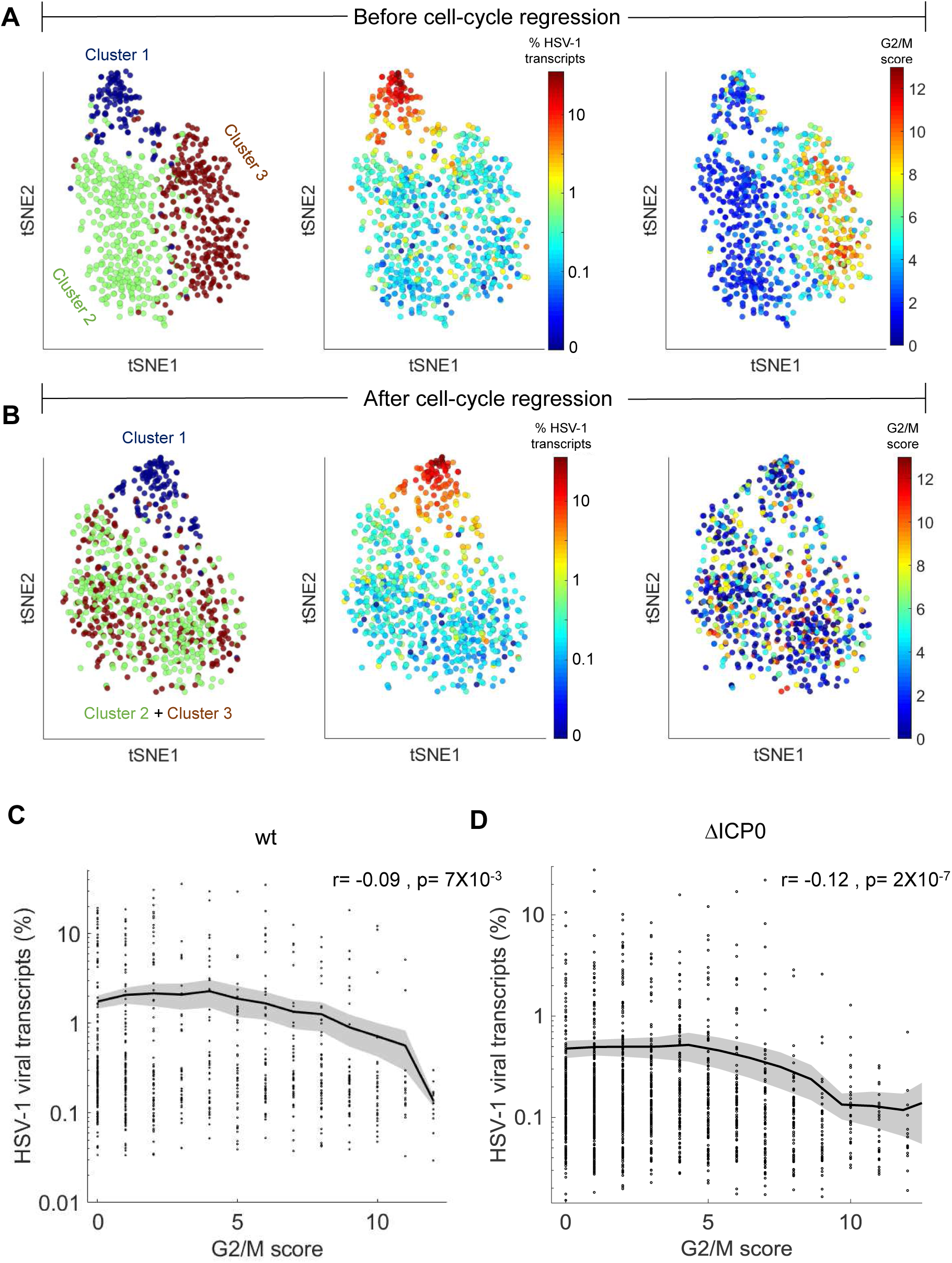
Cell-cycle is anti-correlated with viral gene expression and is a major source of transcriptional variability. **A.** tSNE plots based on viral and host gene expression (wt infection). Left - Cells are colored according to their clustering. Cluster 1 is colored blue, cluster 2 is colored green and cluster 3 is colored brown. Middle - Cells are colored based on the % of viral transcripts they express from blue (low) to red (high). Color bar is logarithmic. Right - Cells are colored based on cell-cycle score, from blue (low) to red (high). . **B.** tSNE plots as in panel A, showing cell clustering (left), viral gene expression (middle) and cell-cycle score (right) after regressing out the cell-cycle effect on gene expression. **C,D.** Cell-cycle score is anti-correlated with wt (C) and ΔICP0 (D) viral transcription. r and p are the Pearson correlation coefficient and associated p-value, respectively

**Supplementary Figure 3.**
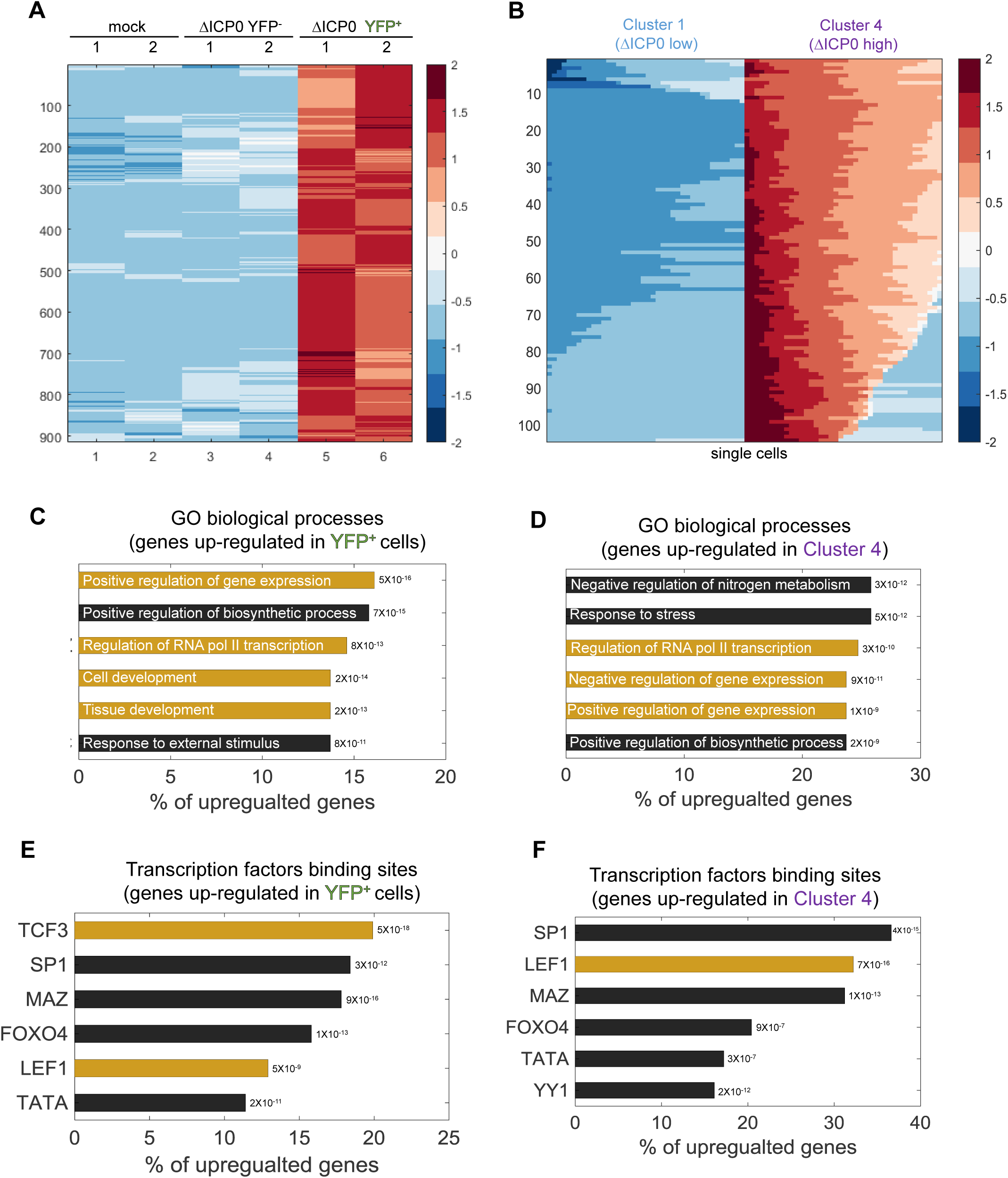
ΔICP0 infection up-regulates developmental pathways. **A.** Heat-map of genes which are significantly up-regulated in ICP4^+^ cells, as compared to both mock and ICP4^-^ cells. RNA-sequencing was performed in duplicates denoted by the numbers 1 and 2 on the top row. Each row shows the normalized (z-score) expression of a single gene, colored from low (blue) to high (red). **B.** Heat-map of genes which are significantly up-regulated in cluster 4 (highly-infected) compared to cluster 1 (lowly-infected) single cell. Each row shows the normalized (z-score) expression of a single gene, colored from low (blue) to high (red) in 80 single cells (40 from cluster 4 and 40 from cluster 1). **C,D.** Go terms enriched for genes identified in panels A and B, respectively. The GO term is written on the bar and the value next to each bar is the FDR-corrected p-value for its enrichment. Bar length shows the % of up-regulated genes annotated to the GO term. Highlighted in yellow are GO terms associated with development and gene regulation. **E,F.** Transcription factors binding sites enriched in the promoters of genes identified in panels A and B, respectively. The value next to each bar is the FDR-corrected p-value for its enrichment. Bar length shows the % of up-regulated genes containing binding sites for each transcription factor. Highlighted in yellow are the LEF1 and TCF3 transcription factors, which are part of the WNT/β-catenin pathway.

**Supplementary Figure 4.**
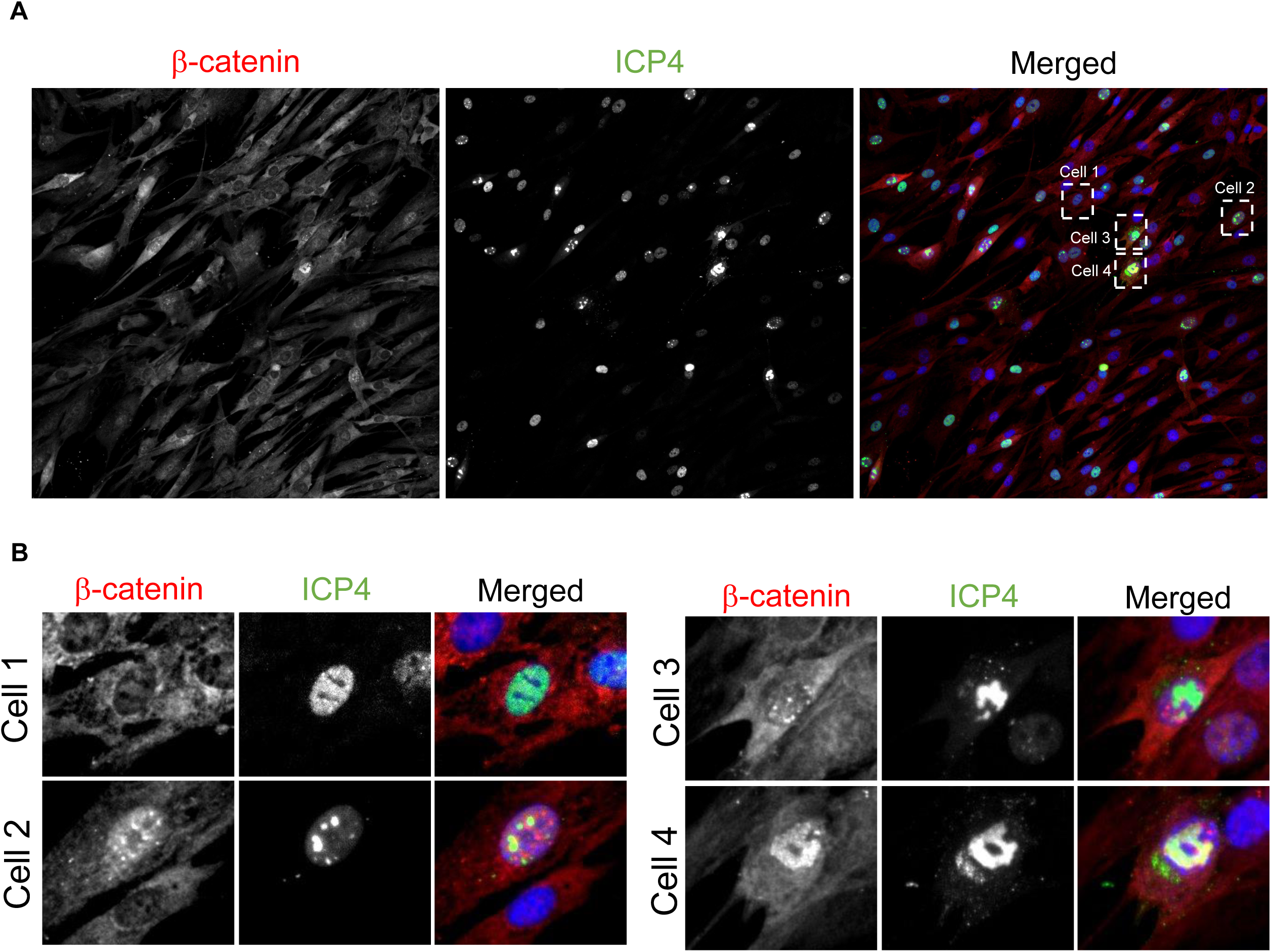
β-catenin translocates to the nucleus and concentrates in the viral replication compartments upon ΔICP0 infection. **A.** Immunofluorescent staining of β-catenin in ΔICP0 infected HDFn cells at 5 hours post infection. **B.** magnified imaged of the four cells denoted by dashed white boxes in panel A, showing representative images of cells with different ICP4 and β-catenin localizations. Note that ΔICP0 infection results in more abortive infections than wt infection, manifesting in many ICP4+ cells which show diffused nuclear localization, unable to form replication compartments.

## MATERIALS AND METHODS

### Cells, viruses and inhibitors

Primary neonatal human dermal fibroblast (HDFn) were purchased from Cascade Biologics (cat #C0045C), grown and maintained in medium 106 (Cascade Biologics, cat #M106500) supplemental with Low Serum Growth Supplement (Cascade Biologics, cat #S00310). Cells were maintained for up to eight passages, and experiments were performed on cells between passages 4-7. A549 cells were purchased from Sigma-Aldrich and maintained in DMEM supplemented with 10% fetal bovine serum. Mel624, a patient-derived melanoma cell-line, was obtained from the lab of Professor Thomas Gajewski at the Univeristy of Chicago and maintained in RPMI supplemented with HEPES, NEAA, Pen/Strep and 10% fetal bovine serum. Vero and U2OS cells (obtained from the lab of Matthew D. Weitzman, University of Pennsylvania) grown in DMEM supplemented with 10% fetal bovine serum were used for viral propagation and titration. wt and ΔICP0 HSV-1 (strain 17) expressing ICP4-YFP were generated by Roger Everett (Everett et al., 2003) and were a kind gift from Matthew D. Weitzman. Viral stock was prepared by infecting Vero cells (for wt virus) or U2OS cells (for ΔICP0) at an MOI of 0.01 and harvesting viral progeny 2-3 days later using 3 cycles of freezing and thawing. Viral stock was titrated by plaque assay, aliquoted and stored at −80C. iCRT14, a β- catenin inhibitor was purchased from Sigma-Aldrich (cat #SML0203) and dissolved in DMSO.

### Time lapse fluorescent imaging

HDFn cells were seeded on 6-well plates and allowed to attach and grow for one day. On the day of the experiment, cells were counted and infected with HSV-1 at an MOI of 2. Cells were washed once with 106 medium without supplements and virus was added in the same media at a final volume of 300μl per well. Virus was allowed to adsorb to cells for one hour at 37C with occasional agitation to avoid cell drying. The inoculum was aspirated and 2ml of full growth media was added and this point was considered as “time zero.” Cells were imaged in a Nikon Ti-Eclipse, which was equipped with a humidity and temperature control chamber. Images were acquired every 15 minutes for 24 hours from multiple fields of view. Image analysis was performed with ImageJ and MATLAB.

### Single-cell RNA-sequencing

HDFn infected with HSV-1 at an MOI of 2 were harvested at 5 hours post-infection and washed three times in PBS containing 0.01% BSA. Cells were counted and processed according to the Drop-seq protocol (Macosko et al., 2015) in the Genomics facility core at the University of Chicago. Sequencing was performed on the Illumina NextSeq500 platform. Preliminary data analysis (quality control, trimming of adaptor sequences, UMI and cell barcode extraction) was performed on a Linux platform using the Drop-seq Tools (Version 1.13) and Drop-seq Alignment Cookbook (Version 1.2), available at https://github.com/broadinstitute/Drop-seq/releases. Alignment of reads was performed using the STAR aligner (Version 2.5.4b, (Dobin et al., 2013) to a concatenated version of the human (GRCh38 primary assembly, Gencode release 27) and HSV-1 genomes (Genbank accession: JN555585). The HSV-1 genome annotation file was kindly provided by Moriah Szpara (Penn State University). Following the generation of the DGE (digital gene expression) file, further analyses were performed in MATLAB and included quality control, cell clustering, correlation and differential gene expression analyses and data visualization. All the scripts used for data analysis and visualization are available upon request.

### RNA-sequencing of sorted cells

HDFn cells were mock or HSV-1 infected at an MOI of 2, trypsinized, washed and re-suspended in full growth media. Cells were filtered through a 100μm mesh into FACS sorting tubes and kept on ice. HSV-1 infected cells were sorted into two populations based on ICP4-YFP expression. 0.5 million cells were collected from each population. Mock-infected cells were similarly sorted. ICP4 negative cells had the same level of YFP fluorescence at mock-infected cells. For ICP4 positive cells, we collected cells that were in the top 30% of YFP expression. The two populations were clearly separated from each other. Sorting was performed on an AriaFusion FACS machine (BD) at the University of Chicago flow-cytometry core facility. Total RNA was extracted from cells using the RNeasy Plus Mini Kit (QIAGEN) and submitted to The University of Chicago Genomics core for library preparation and sequencing on a HiSeq4000 platform (Illumina). Reads were mapped to a concatenated version of the human and HSV-1 genomes with STAR aligner (see single-cell RNA-sequencing above for details). Reads were counted using the featureCounts command, which is a part of the Subread package (Liao et al., 2013). Further analyses were performed in MATLAB and included differential gene expression analyses and data visualization. All the scripts used for data analysis and visualization are available upon request.

### Sequencing data availability

All sequencing data has been deposited in the Gene Expression Omnibus (GEO) under accession number 396 GSE126042.

### Immunofluorescence staining

HDFn were seeded in 24-well plates and allowed to attach and grow for one day. Cells were infected as above and fixed using a 4% paraformaldehyde solution at 5 hours post-infection. Cells were fixed for 15 minutes at room temperature and washed, blocked and permeabilized with a 10% BSA, 0.5% Triton-X solution in PBS for one hour. Cells were then incubated with primary antibodies in a staining solution (2% BSA, 0.1% Triton-X in PBS) overnight at 4C. Cells were washed three times with PBS, incubated with secondary antibodies in staining solution for 1 hour at room temperature, washed three times with PBS and covered with 1 ml PBS containing a 1:10,000 dilution of Hoechst 33342 (Invitrogen, cat #H3570). Cells were imaged on a Nikon Ti-Eclipse inverted epi-fluorescent microscope. Primary antibodies were mouse monoclonal anti-β-catenin (R&D systems, cat #MAB13291, used at 1:200 dilution) and rabbit monoclonal anti-IRF3 (Cell Signaling Technologies, Cat #11904S, used at 1:400 dilution). Secondary antibodies were AlexaFluor 555 conjugated anti-mouse and anti-rabbit F(ab’)2 fragments (Cell Signaling Technologies, cat 409 #4409S, #4413S, used at 1:1,000 dilution).

### siRNA nucleofection

5×10^5^ HDFn were washed once in PBS and nucleofected with 1µM siRNA against β-catenin (Dharmacon, siGENOME Human CTNNB1, cat #M-003482-00-0005) or a scarmbeld siRNA control (Dharmacon siGENOME Non-Targeting siRNA Pool #1, cat #D-001206-13-05) using the Human Dermal Fibroblast Nucleofector^TM^ Kit (Lonza, cat #VPD-1001). β-catenin expression was assayed 3 days later by Q-PCR.

## ACKNOWLEDGMENTS

We wish to thank Matthew D Weitzman for sharing with us the ICP4-YFP expressing HSV-1 and Moriah Szpara for the genome annotation of the HSV-1 strain 17. Sequencing and library preparations were performed at The University of Chicago Genomics core facility and cell sorting at the Flow Cytometry core. We also with to thank Oren Kobiler for support and advice throughout the project. ND wishes to thank EMBO and HFSPO for their support through post-doctoral fellowships at different stages of the project.

## AUTHOR CONTRIBUTIONS

ND and ST designed the experiments, ND, PP and LV performed experiments, ND preformed data analysis, ND and ST wrote the manuscript.

## DECLARATION OF INTERESTS

No conflicts of interests exist

